# EEG Responses to Upper Limb Pinprick Stimulation in Acute and Early Subacute Motor and Sensorimotor Stroke

**DOI:** 10.1101/2024.06.05.597652

**Authors:** L Tedesco Triccas, S Van Hoornweder, TC Camilleri, L Boccuni, A Peeters, V Van Pesch, R Meesen, D Mantini, KP Camilleri, G Verheyden

## Abstract

**Background:** Electroencephalography (EEG) during pinprick stimulation has the potential to unveil neural mechanisms underlying sensorimotor impairments post-stroke. This study explored event-related peak pinprick amplitude and oscillatory responses in healthy controls, in people with motor and sensorimotor in acute and early subacute stroke, their relationship and to what extent EEG somatosensory responses can predict sensorimotor impairment.

**Methods:** In this study, involving 26 individuals, 10 people with a (sub-)acute sensorimotor stroke, 6 people with a (sub)acute motor stroke and 10 age-matched controls, pinpricks were applied to the dorsa of the impaired hand to collect somatosensory evoked potentials. Time(-frequency) analyses of somatosensory evoked potential (SEP) data at electrodes C3 and C4 explored peak pinprick amplitude and oscillatory responses across the three groups. Also, in stroke, (sensori-)motor impairments were assessed at baseline Fugl Meyer Assessment Upper Extremity (FMA-UE) and Erasmus modified Nottingham Sensory Assessment (EmNSA) at baseline and 7 to 14 days later including Fugl Meyer Assessment Upper Extremity (FMA-UE) and Erasmus modified Nottingham Sensory Assessment (EmNSA). Mixed model analyses were used to address objectives.

**Results:** It was demonstrated that increased beta desynchronization magnitude correlated with milder motor impairments (R^2^ =0.213), whereas increased beta resynchronization and delta power were associated to milder somatosensory impairment (R^2^ =0.550). At the second session, larger peak-to-peak SEP amplitude and beta band resynchronization at baseline were related to greater improvements in EMNSA and FMA-UE score, respectively, in sensorimotor stroke group.

**Conclusions:** These findings highlight the potential of EEG combined with somatosensory stimuli to differentiate between sensorimotor and motor impairments in stroke, offering preliminary insights into both diagnostic and prognostic aspects of upper limb recovery.

## Background

Somatosensory integration plays a critical role in precise motor control and body awareness (1). Despite its importance, upper limb stroke research predominantly focusses on motor impairments and their impact on recovery (2). However, in the first month post-stroke, a significant number of individuals –ranging from 21 to 54% of people with stroke– experience somatosensory, which can impede recovery (3-7). The contribution of somatosensory factor to upper limb sensorimotor impairment in the early-stage post-stroke remains poorly understood.

To an extent, this lack of comprehension may stem from limitations of currently used measures of somatosensory function, particularly their lack of precision and suitability for people with aphasia. Notably, these limitations may, in part, explain previous non-significant results in research assessing the impact of upper limb somatosensory rehabilitation programmes (8-10).

Electroencephalography (EEG) presents a promising avenue to address some of these shortcomings by providing a portable, inexpensive, and non-invasive method to register brain activity on the scalp. Indeed, Stroke Recovery and Rehabilitation Round tables and recent reviews have recommended exploring EEG measures as biomarkers for stroke recovery, aiming to improve our understanding of brain dynamics in stroke (11-13). When combined with somatosensory evoked potentials (SEPs), EEG can provide information on the integrity of the central nervous system’s motor and somatosensory pathways. Notably, the absence of SEPs indicates poor upper limb recovery (14-16).

Although limited, previous work investigating SEPs in stroke has proven insightful. For one, reduced amplitudes and increased latencies of SEPs generated by median nerve stimulation have been associated with poorer upper limb motor function of the upper limb in acute and early subacute stroke (17, 18). Abnormal median nerve SEP components have also been identified in people with acute pure sensorimotor stroke, with N_2_ and P_2_ potentials being absent or decreased in amplitude (19). Yet, the predictive value of SEPs for sensorimotor recovery remains unclear.

Oscillatory activity registered by EEG in the delta, theta, alpha and beta bands has also been proposed as a potential for upper limb motor recovery (20-23). For instance, decreased alpha power has been proposed as potential biomarker for upper limb motor recovery (20-22). Likewise, a decreased beta band rebound during passive movement has also been reported in persons with pure upper limb motor stroke, with rebound magnitude demonstrating a correlation to hand motor clinical scores (23).

In contrast to median nerve stimulation and passive movements, pinprick stimulation yields the advantages of being a pure somatosensory stimulus that is easy to combine with EEG, and well-controlled in terms of stimulus intensity (24-26). Indeed, our recent work in healthy controls highlights that pinprick stimuli on the dorsa of the hands are highly reliable to explore oscillatory activity in the delta, theta and alpha frequency bands in first 0.25s (27).

Currently, a comparison of pinprick SEPs in (early sub-)acute motor stroke, (early sub)acute sensorimotor stroke, and age-matched controls is lacking. However, such a comparison is imperative if SEPs are to be used as biomarkers in stroke, as recommended by previous guidelines. Moreover, the association between SEPs and functional outcome measures in acute motor and sensorimotor stroke remains to be explored. Therefore, we set out to address the following research questions; (1) Is there a difference in event-related peak amplitude and oscillatory EEG data, two common SEP metrics, because of pinprick stimulation, in motor-versus sensorimotor-stroke versus healthy controls? (2) Is there a relationship between SEP measures and clinical somatosensory and motor measures of the upper limb? (3) To what extent can SEP measures predict upper limb sensorimotor recovery in people with (sub-)acute stroke?

Concerning pinprick SEP amplitude analyses, we hypothesised that people with acute stroke to have longer latencies and reduced amplitudes which will correlate with the severity of somatosensory and motor deficits (18, 23). The absence of prior research in acute stroke using time-frequency analyses, prompted us to adopt an exploratory approach.

## Material and Methods

Ethical approval was obtained from the Ethics Committee of the University Hospital of Leuven (S61174) and the University of Malta Research Ethics Committee (002/2016). An overview of the current observational study can be found in **Fig 1**. Data collection commenced on 28^th^ January 2019 and completed on 10^th^ August 2019.

**Fig 1.**
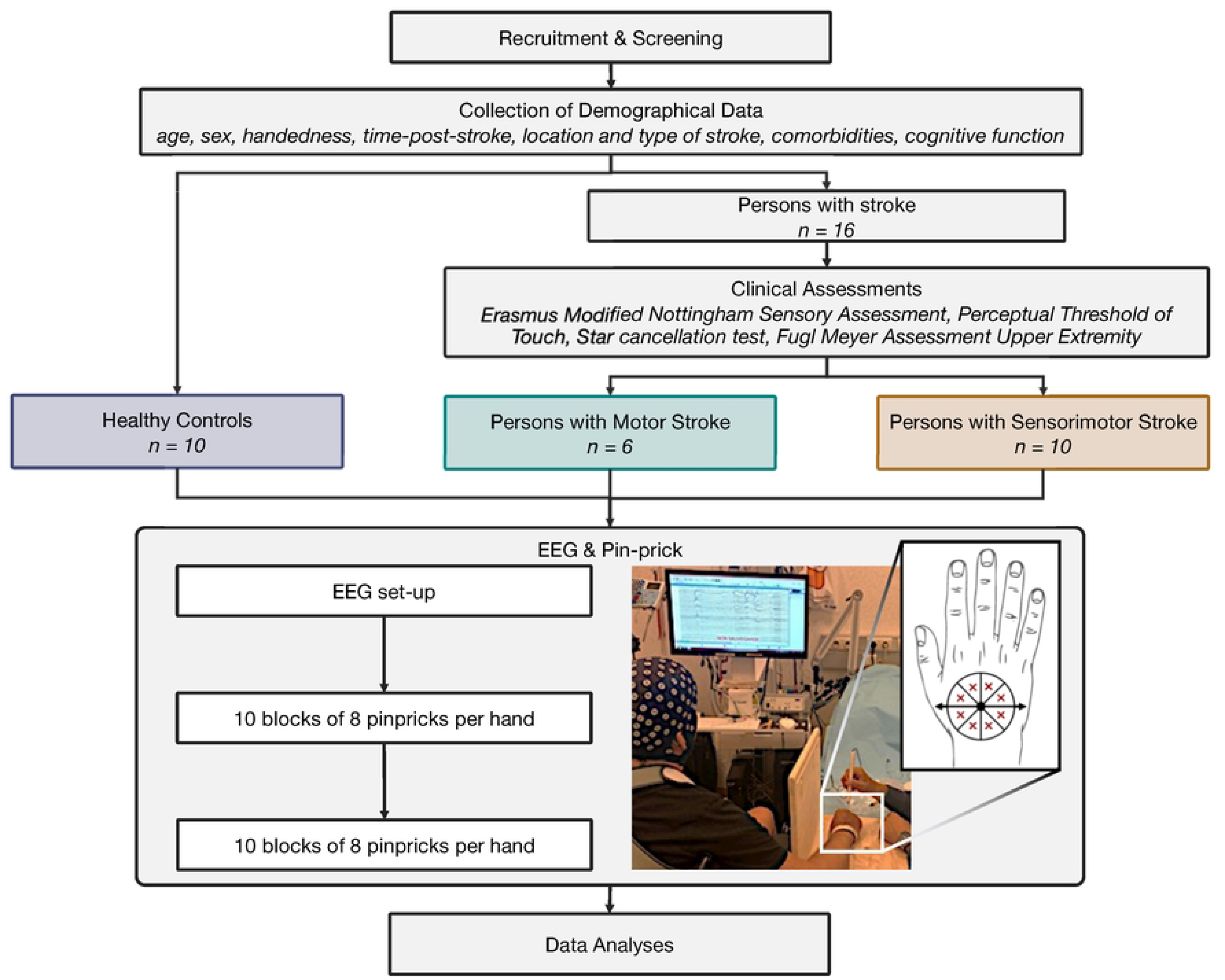
Global overview of study and electroencephalography (EEG) – pinprick protocol.

### Participants

People with stroke were recruited from the stroke unit of the Saint Luc Hospital, Brussels (Belgium) and the Mater Dei Hospital, Msida (Malta). To do so, eligible persons and/or a family member were informed about the study via the ward’s neurologist(s) and/or physiotherapist(s). When interested, they then received a visit by the principal researcher who explained the study in detail and answered all questions. Subsequently, participants gave their written informed consent. Age-matched healthy adults without any somatosensory impairments were recruited in Malta via word of mouth. For persons with stroke, inclusion criteria were: (1) ≥ 18 years old, (2) a first-ever unilateral, supra-tentorial stroke as defined by the World Health Organisation (28), (3) somatosensory upper limb deficits, as indicated by the Rivermead assessment of somatosensory performance (29) and/or motor upper limb deficits, as measured by Fugl Meyer Assessment Upper Extremity (FMA-UE) (30).

### Clinical Assessments

In persons with stroke, clinical assessments were collected at baseline (T1) and 7 to 14 days after the initial study visit (T2): (i) Erasmus modified Nottingham Sensory Assessment (EmNSA), involving measuring tactile sensation (score out of 32) and proprioception (score out of 8) of both upper limbs (31), (ii) Perceptual Threshold of Touch (PTT), determining the smallest possible stimulus intensity a person can detect using a 2-channel TENS CEFAR tempo stimulator with 3 cm self-adhesive electrodes placed on the bulb of the index finger (anode) and the palm (cathode) of the hand (32), (iii) Star cancellation test measuring visuospatial neglect, and containing 52 large stars, 13 letters and 10 short words surrounded by 56 smaller stars, with crossing ≤44 indicating unilateral spatial neglect (33) and (iv) FMA-UE to examine motor impairment (score out of 66).

### Electroencephalography and pinprick stimulation data collection

Following the clinical assessments for participants with stroke and directly for healthy controls, the EEG protocol started by placing electrodes on the head according to the extended 10-20 international system (34). In Malta, the g.tec system with 32 g.Scarabeo sintered Ag-AgCl ring electrodes was used, the ground was placed on AFz and the reference clip was attached to the left ear lobe of the participant. In Belgium, the Micromed Limited system was used with 28 sintered Ag-AgCl electrodes in a cap of ANT NEURO Limited. The ground was placed on POz and the reference clip was attached to the forehead of the participant. Electrode caps were placed to ensure that Cz was exactly in the middle of an imaginary line drawn from the occipital tuberculum posterior to the middle of the nasal bridge. The scalp was prepared by means of a cotton swab, and conductive gel was applied to reduce impedance to ≤ 10 kΩ.

During EEG data acquisition, participants were seated in a (wheel)chair or in supine, lying on their hospital bed. They were asked to sit as still as possible while fixating their gaze and keeping their hand palms facing downwards, resting on a piece of wood, allowing the researcher to apply pinprick stimulation. As shown in **Fig 1**, vision of participants in relation to the tested hand was blocked by means of a screen.

Pinprick application was performed in a standardized manner (27) In both hands, the distance between the base of metacarpal I to the middle of metacarpal V was measured. The middle of this distance served as the centre of a circle drawn on the dorsum of the hand. This circle was divided in eight equal compartments (cf., **Fig 1**). Via a pinprick stimulator (MRC Systems, Germany), sharp stimuli were applied, and triggers were sent to the EEG device. To ensure that all pinpricks were applied with nearly identical force, all researchers involved in data acquisition underwent a training programme. Ten sets of 8 stimuli (one stimulus per compartment) were applied in a random order on the dorsa of both hands in an alternating manner. This procedure was carried out twice, totalling 160 stimuli per hand. EEG data were acquired at a sample rate of 256 Hz.

### Data Analyses

Demographic and clinical assessment data were aggregated in an .xlsx file. EEG data were pre-processed offline in Matlab (Mathworks® v R2018a, Natick, Maine, USA) using custom code based on the EEGLAB plug-in (Swartz Centre for Computational Neuro-science, eeglab 14_1_2b) (35). Specifically, EEG data were 1 – 35 Hz forward-backwards band-pass filtered with a FIR filter, noisy channels were removed via Clean RawData (36). Next, data were re-referenced to the common average reference, bad data periods were removed through Artefact Subspace Rejection, individual component analysis (ICA) was conducted to remove artefactual components in an automated manner through ICFlag.

#### EEG Time-Domain Analyses

For time-domain analyses, EEG epochs were extracted with a window of -0.5 s to 1.5 s, with 0s being pinprick administration. EEG epochs were averaged across trials, separately for each hand and condition. Epochs were baseline-corrected with a reference interval of -500 to 0 ms. For each average epoch, the minimum and maximum amplitudes and corresponding latencies were extracted in the channels C3 and C4.

#### EEG Time Frequency Analyses

For the time-frequency analyses, a similar approach as Van Hoornweder et al (37) was used which implemented custom code based on the Cohen (2014) (38). Epochs from -1.5 s to 2.2 s were extracted from the pre-processed EEG data, with 0 s being pinprick administration. Data within these epochs from channels C3 and C4 were convoluted with complex Morlet wavelets, constructed as Gaussian-windowed complex sine waves:

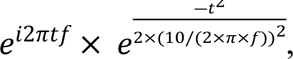

where *i* = imaginary operator, *t* = time, and *f* = frequency going from 1 to 35 Hz in 30 logarithmic steps. From this complex signal, frequency-specific power values were extracted at each time point using the squared magnitude of the result of the convolution. Power values obtained from the time-frequency decomposition were dB normalized, with baseline being the frequency-specific average power values from −0.4 s to −0.1 s.

Subsequently, we extracted frequency specific power values per participant, channel and condition using a frequency and temporally indeterministic bootstrapping-based masking procedure (**Fig 2**). First, we computed a time-frequency matrix averaging together all activity across all three groups and all pinpricks, mitigating the risk of circular interference influencing the mask and our subsequent results.

**Fig 2.**
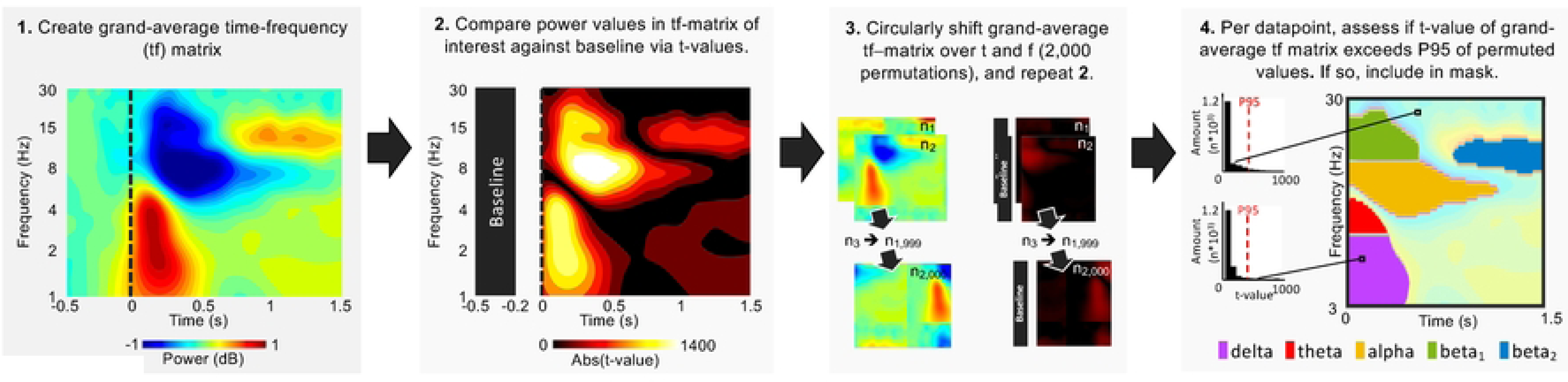
Frequency and temporally indeterministic bootstrapping-based masking procedure used to identify time-frequency (tf) regions of interest.

Second, each datapoint from 0 to 1.6 s was tested against the baseline period activity via a one-sample T-test, retrieving a t-value per datapoint. Third, the grand-average matrix obtained in step 1 was circularly shifted over time and frequency and step 2 was repeated for the obtained matrices. This third step was reiterated 2,000 times, resulting in a null-distribution of 2,000 t-values per data point in the time-frequency matrix. Fourth and finally, per data point, we assessed whether the t-value of the grand-average time-frequency matrix (step 1) was in the ≥ 95^th^ percentile of the t-values obtained by the permutation procedure (step 3). If so, the time-frequency point was included in the time-frequency mask.

The results of this procedure, shown in **Fig 2, panel 4**, were five regions of interest: (1) delta band synchronization (1 – 4 Hz, 0 ms to 543 ms), theta band synchronization (4 – 6 Hz, 0 ms to 262 ms), alpha band desynchronization (5 – 13 Hz, 47 ms to 785 ms), beta band desynchronization (13 – 27 Hz, 70 ms to 566 ms) and beta band resynchronization (11 – 17 Hz, 820 ms to 1601 ms). In persons with stroke, mean power was extracted from these regions of interest from the affected hemisphere, when the pinprick was applied to the affected limb. In healthy persons, mean power was extracted from these regions of interest in the time-frequency data from the hemisphere contralateral to where each pinprick was applied.

### Statistical Analysis

To assess whether the healthy controls were age-matched, a one-way ANOVA was applied to the data in IBM SPSS Statistics 28. All other statistical analyses were conducted in RStudio (39), via lmerTest (40), ggplot2 (41), Emmeans (42). For all tests, alpha was set to 0.05.

To investigate if the (dis)similarity of SEPs in healthy controls, persons with motor stroke and persons with sensorimotor stroke, amplitude and time-frequency properties were statistically analyzed. When relevant, significant fixed effects were subjected to Tukey-corrected post-hoc tests. If significant differences were present, Cohen’s d effect sizes were extracted.

Concerning the EEG ERP amplitude data, a one-way ANOVA was established with AMPLITUDE as dependent variable. The fixed effect was GROUP (healthy, motor stroke, sensorimotor stroke).

For the EEG time-frequency data, a linear mixed effects model was established with POWER as dependent variable. PARTICPANT was included as random intercept. The fixed effects were GROUP (healthy, motor stroke, sensorimotor stroke), TIME-FREQUENCY FEATURE (delta synchronization, theta synchronization, alpha desynchronization, beta desynchronization, beta resynchronization), and GROUP * TIME-FREQUENCY FEATURE. Stepwise backward model building was performed to obtain the most parsimonious model.

To unveil the association between the various EEG features and clinical measurements, two linear regression analyses were performed in the stroke cohort for the three clinical scores that were of interest; the FMA, EMNSA and PPT. Akin to the EEG analyses, we focused on the clinical scores of the affected side. The first linear regression analyses investigated the effect of the TIME-and TIME-FREQUENCY EEG FEATURES (peak-to-peak amplitude, delta, theta, alpha, beta1 and beta2), together with STROKE TYPE (motor or sensorimotor), on the clinical scores. Beyond the main effects, the interaction between each EEG FEATURE * STROKE TYPE was also analysed. The second linear regression model yielded the same fixed effects, but the dependent variable was the change in clinical score from baseline (T1) to 7 to 14 days after the initial measurement (T2). Thus, this second model informed on the predictive value of the EEG time(-frequency) features as a short-term predictive marker. Here also, stepwise backward model building was used.

## Results

### Demographics and clinical profiles of the participants

Sixteen participants with stroke (sensorimotor group (n=10): mean age: 63.7 ±9.53; motor group (n=6): mean age: 73.2 ±11.3), and ten age-matched healthy controls (mean age: 62.3±10.3) were included. While absence of evidence is not evidence for absence, the ANOVA investigating if age was significantly differed across the three groups did not reject the null hypothesis (p-value = 0.133) (**Table 1**). Clinical profiles of both the sensorimotor and motor stroke groups are presented in **Table 1**. No participant presented with neglect. Impairments in No participant presented with neglect. Impairments in touch were present in the sensorimotor group, as indicated by the EMNSA and the PTT. Both the motor and sensorimotor stroke groups showed moderate motor impairments (43). After analysis, the mean ± standard deviation number of epochs for the stroke groups were n = 110 ± 38 and for the healthy controls was n = 151 ± 20.

**Table 1:**
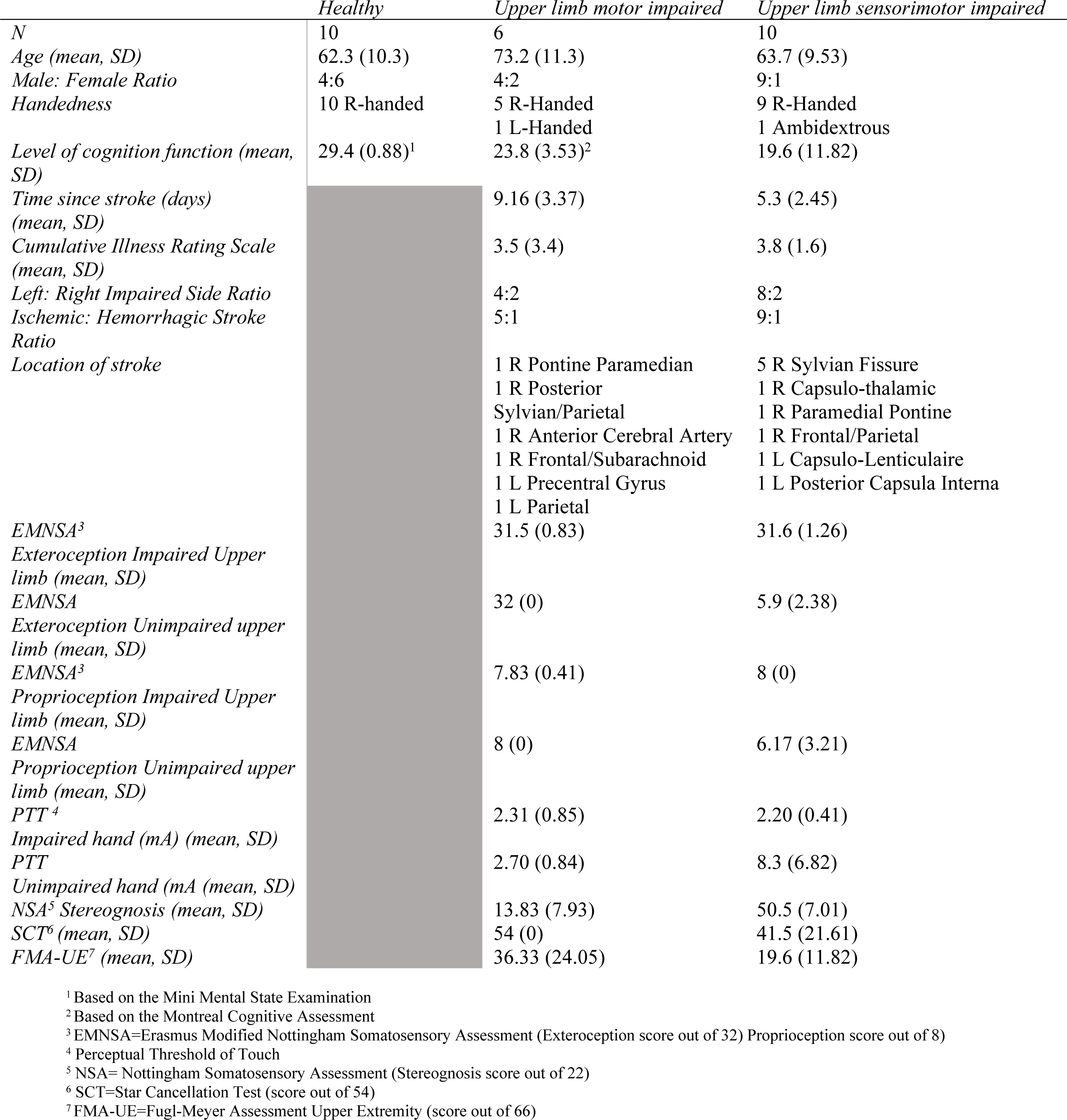
Demographics and clinical characteristics of the sample.

### Time-domain Analysis

The NP amplitude and positive peak latency of the SEP of the dominant healthy hand was 2.81 ± 1.30 µV and 202.69 ± 70.62 ms, the motor impaired was 3.22 ± 2.68 µV and 169.09 ± 76.05 ms and sensorimotor hand was 1.36 ± 0.54 µV and 161.46 ± 66.63 ms. Data from one individual with motor stroke were identified as an outlier based on the normal quantile plots and removed to adhere to the assumption of normality.

The one-way ANOVA investigating the peak-to-peak amplitude of time-domain EEG activity in the central region contralateral to the pinprick, did not contain an effect of GROUP (F_2_ = 3.217, p = 0.055) (**Fig 3**). While the observed effect did not reach significance, the exploratory nature of our current study, prompted us to conduct Tukey-corrected post-hoc tests.

**Fig 3.**
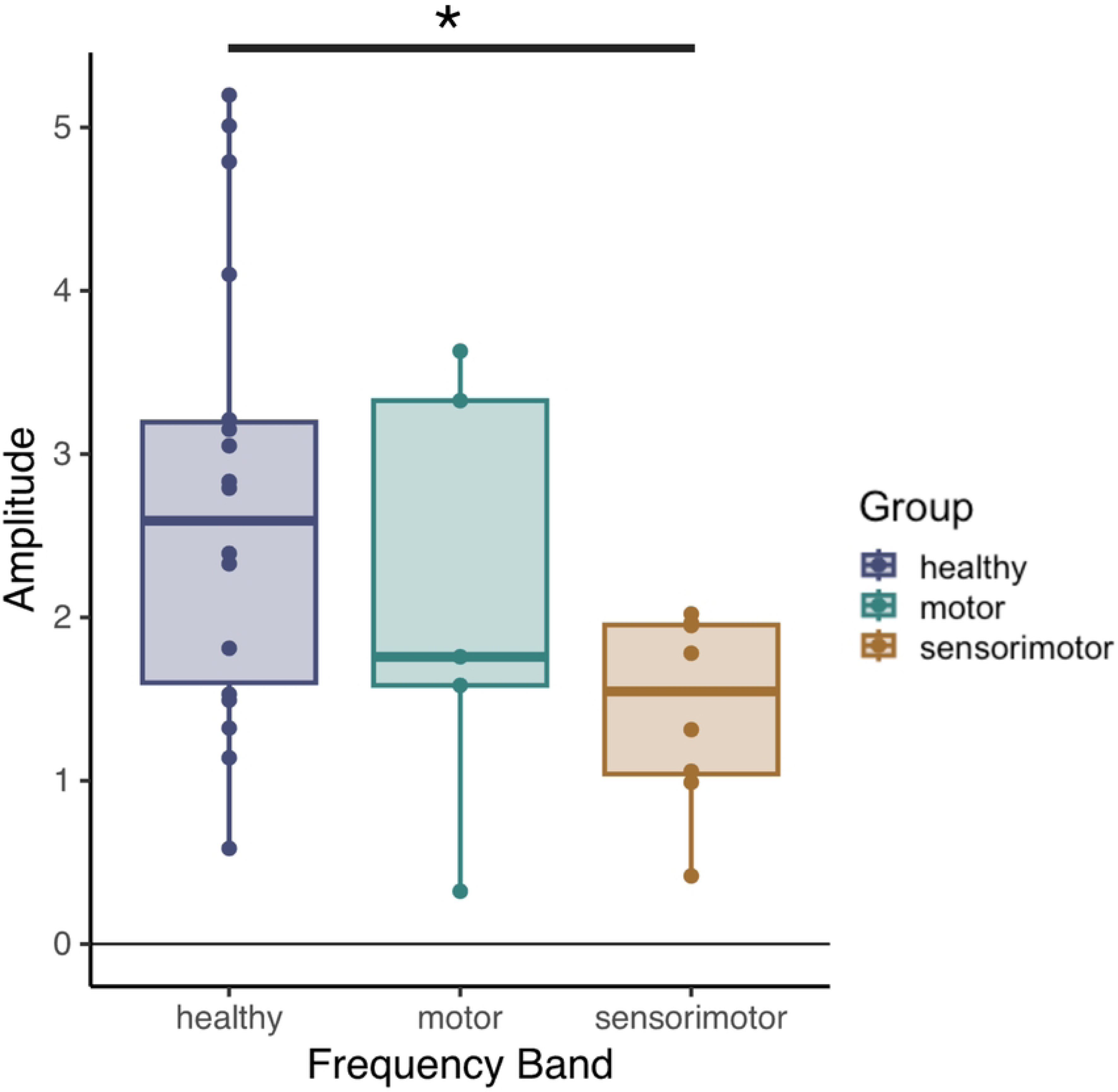
Pin prick evoked potential amplitude for three groups; healthy, motor and sensorimotor

Post-hoc tests found a large difference in peak-to-peak amplitude of the pinprick SEP in healthy controls (peak-to-peak amplitude = 2.73 ± 1.36 µV, mean ± standard deviation), compared to persons with stroke in the sensorimotor group (peak-to-peak amplitude = 1.44 ± 0.59 µV) (t_28_ = 2.512, p = 0.046, Cohen’s d = 1.067). There were no significant differences between healthy controls and people with stroke in the motor group (peak-to-peak amplitude = 2.12 ± 1.36 µV) (p = 0.590, Cohen’s d = 0.499), and in the motor compared to the sensorimotor groups (p = 0.585, Cohen’s d = 0.568).

### Time-frequency Analysis

One outlier datapoint was removed to ensure normality (delta band power of a person with motor stroke). The linear mixed model, investigating the time-frequency activity in the central region contralateral to the pinprick, contained a significant GROUP * FREQUENCY BAND interaction (F_8, 134.4_ = 2.244, p = 0.028). Tukey-corrected post-hoc tests were used to interpret this effect. While it seems that this interaction was driven by alpha band desynchronization, which was attenuated in participants with sensorimotor stroke compared to healthy controls, its effect did not survive multiple comparison correction (t_131_ = -2.365, corrected p = 0.051, Cohen’s d = -0.987). Likewise, no other significant effects remained after correction (**Fig 4**).

**Figure 4.**
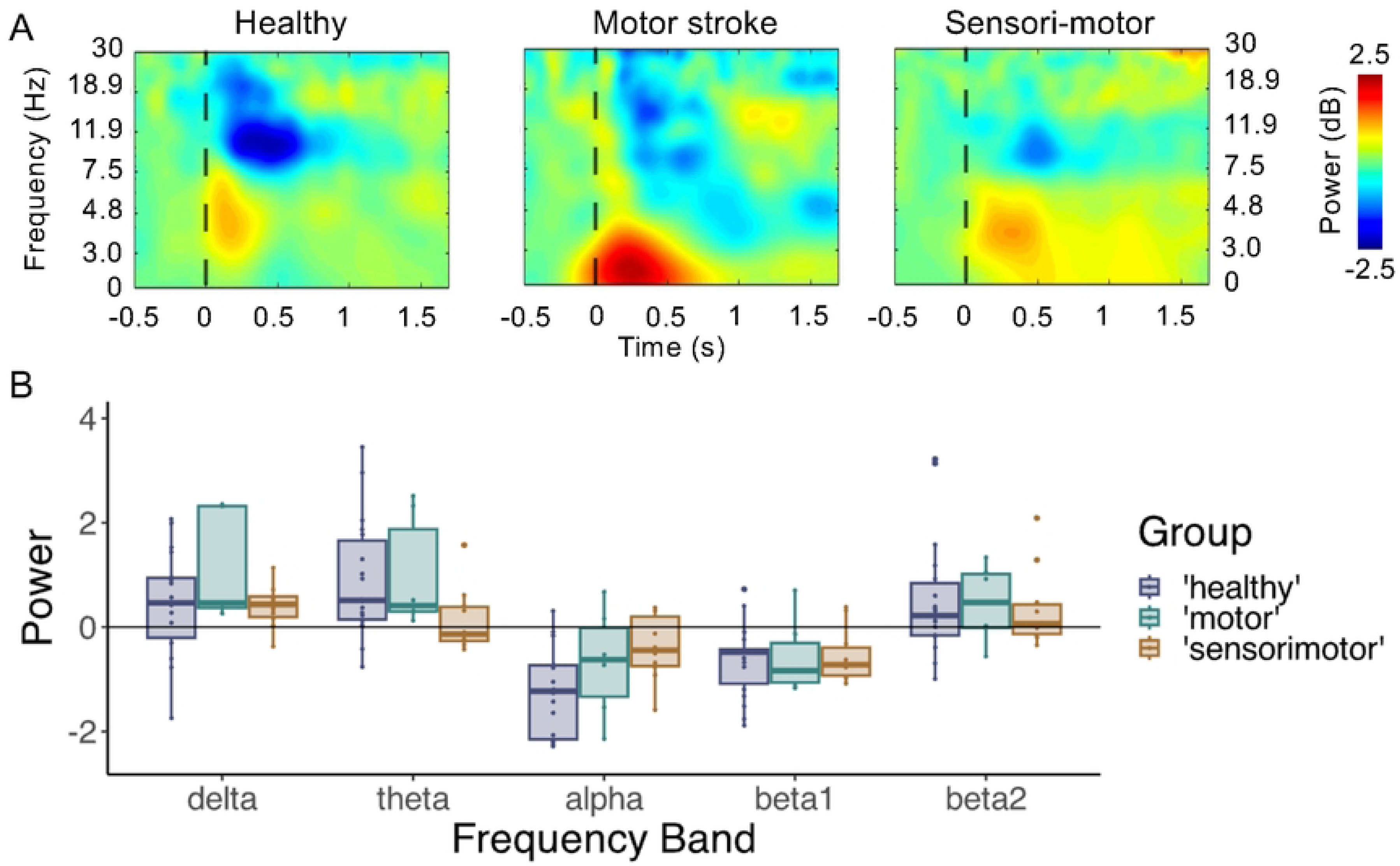
Time-frequency SEPs due to pinprick in participants with stroke with motor and sensorimotor deficits, and healthy controls. **A)** Average time-frequency matrices with 0 s being the application of the pinprick. **B)** Boxplots showing the power from pinprick stimulation in the different time-frequency features of interest. While a significant interaction effect of Frequency Band * Group was present (p = 0.028), no pairwise comparisons survived multiple comparison correction.

### Relationship between event-related time and time-frequency EEG features and clinical somatosensory and motor measures

First, we investigated the link between the EEG features and clinical scores obtained during the same visit.

Concerning the FMA, we withheld a significant effect of beta1 power (i.e., beta desynchronization) (p = 0.041, estimate = -19.26, R^2^ = 0.213) on the initial FMA score. No other EEG features, nor the effect of stroke type remained in the final model. As shown in **Fig 5** (left panel), a higher degree of beta desynchronization (i.e., lower absolute beta1 power) was associated with a higher FMA score.

**Fig 5.**
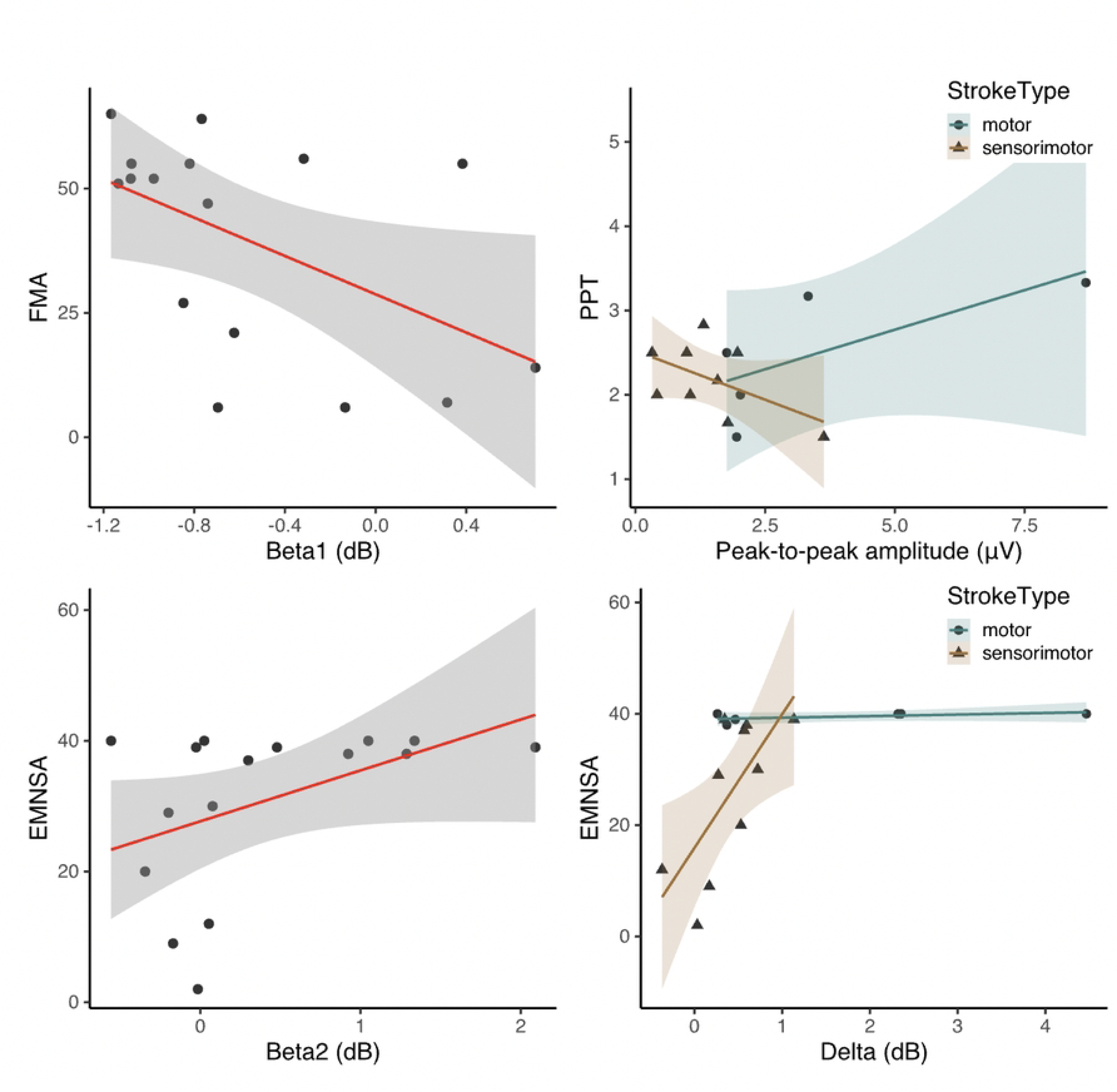
Relationship between EEG features of interest and clinical measures. Upper left panel: A significant effect of beta1 power (i.e., beta desynchronization) on FMA score was found, indicating that a higher degree of desynchronization was related to a better FMA score. Upper right panel: The relation between peak-to-peak amplitude and the PPT score depended on the stroke type, with participants with motor stroke showing a positive relationship whereas the opposite was verified for sensorimotor stroke group. Lower panels: In people with sensorimotor stroke, the amount of delta band synchronization was positively associated to the EMNSA score. In people with motor stroke, this was not the case due to all participants scoring high on the EMNSA score. The amount of beta band resynchronization (i.e. beta2) was positively associated to EMNSA score.

Concerning the PPT, a significant interaction between stroke type * peak-to-peak ERP amplitude was found (estimate = -0.42, p = 0.050, R^2^ = 0.303). As shown in **Fig 5** (right panel), in persons with sensorimotor stroke, a larger peak-to-peak amplitude was associated with lower PPT scores whereas the opposite yielded true for persons with a motor stroke.

Concerning the EMNSA, we observed significant effects of beta 2 power (i.e., beta resynchronization) (estimate = 2.58, p = 0.026) and the interaction between stroke type * delta power (estimate = 21.28, p = 0.025), implying that the main effects of stroke type (estimate = - 22.68, p = 0.006) and delta power (estimate = -0.34, p = 0.894) also were present in the final model (R^2^ = 0.550). As shown in **Fig 5** (lower panels), more beta resynchronization was related to a higher EMNSA score. The interaction between delta power and stroke type was driven by persons with sensorimotor stroke. In this subgroup, delta power and EMNSA score where positively associated. Conversely, a relationship between delta power and EMNSA score was absent in persons with motor stroke, due to all these participants having (nearly)maximal EMNSA scores.

We also assessed whether the improvement in clinical scores from baseline (T1) to 7 to 14 days later (T2) was related to the EEG time(-frequency) features.

For the FMA, the final model contained beta2 (i.e., beta synchronization) (estimate = - 8.96, p = 0.041, R^2^ = 0.214) (**Fig 6**, left panel). A smaller amount of beta synchronization in the first session was associated to a larger improvement in FMA in the second session.

**Fig 6.**
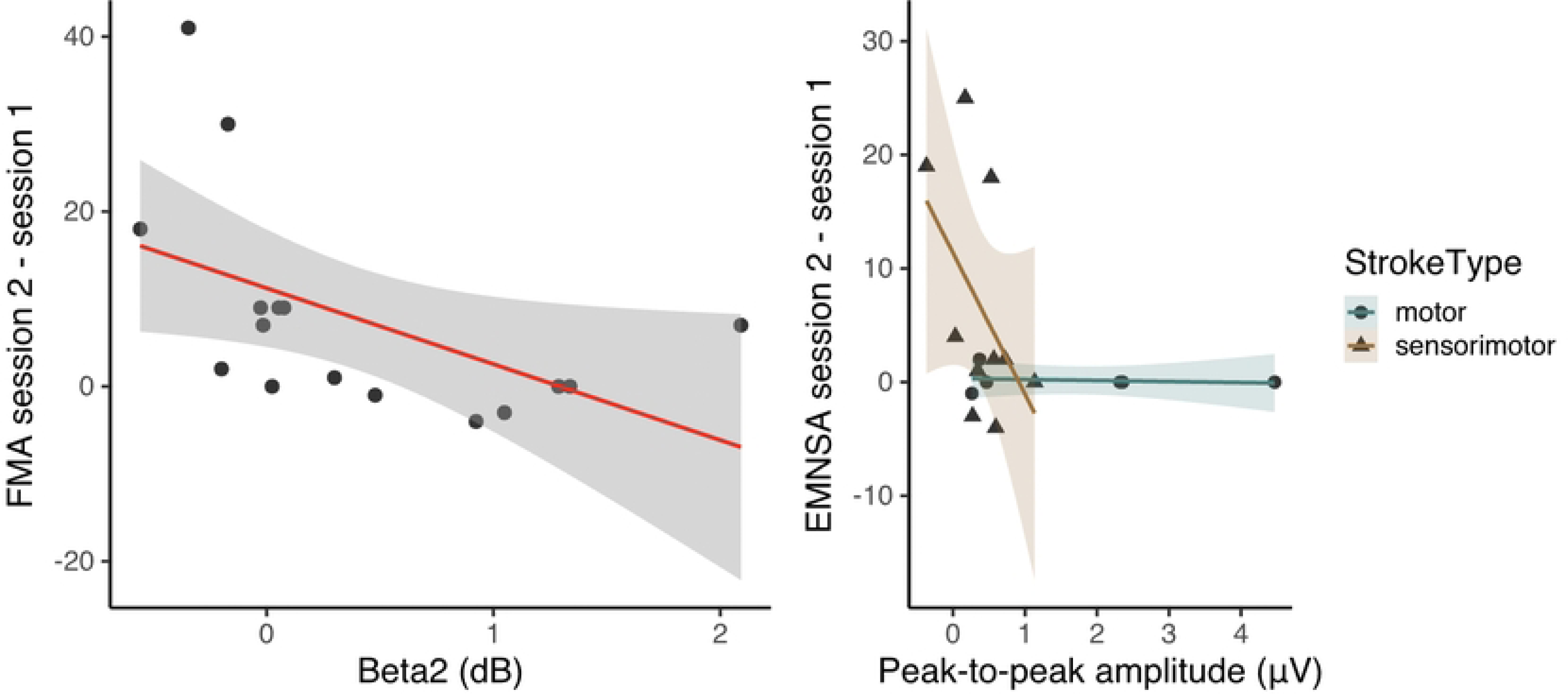
Relationship between EEG time(-frequency) features during session 1 and type of stroke and change in clinical scores from session 1 to session 2 (7 to 14 days later). Left: Low beta resynchronization (i.e., beta2 power) during the initial session was related to a greater improvement in FMA score. Right: Only in people with sensorimotor stroke, the initial peak-to-peak amplitude following the pinprick was positively associated to the degree of improvement in the EMNSA. Due to persons with motor impairments already having high scores during the initial visit, this effect was not present.

Concerning the PPT, the final model contained no fixed effects related to EEG features, only retaining the effect of Stroke Type (estimate = 3.42, p = 0.013, R^2^ = 0.320) (**Fig 6**, middle panel). As expected, participants with sensorimotor stroke showed greater changes in PPT scores 7 to 14 days later compared to motor stroke group, who had intact somatosensory function.

Lastly, for the EMNSA, the final model yielded a significant interaction between stroke type and ERP peak-to-peak amplitude (estimate = 6.61, p = 0.030, R^2^ = 0.332) (**Fig 6**). In sensorimotor stroke, a larger peak-to-peak amplitude during the initial session was related to a greater improvement in EMNSA score. For the people with motor stroke, this effect was not present due to their initial high score (**Fig 5**).

## Discussion

This exploratory study aimed to explore pinprick SEPs in people with acute and early subacute stroke, their relationship with somatosensory and motor clinical measures and their ability to predict short-term upper limb sensorimotor recovery. Our findings underscore the value of EEG features in the context of early-stage upper limb impairment post-stroke. We observed that people with sensorimotor impairments had a significantly smaller negative-positive peak-to-peak SEP amplitude, compared to healthy and purely motor impaired in the first two weeks post-stroke. Also, in the sensorimotor stroke group, SEP peak-to-peak amplitude was associated with somatosensory impairment, with a larger amplitude at baseline being indicative of greater improvement at from baseline to T2, 7 to 14 days later. Abnormal delta frequency band power was related to somatosensory impairments measured by the EMNSA. Across both stroke types, increased beta desynchronization and resynchronization were also related to milder motor and somatosensory impairments, respectively. Also, larger peak-to-peak SEP amplitude and beta band resynchronization at baseline were related to greater improvements in EMNSA and FMA-UE scores, respectively, in sensorimotor stroke.

Previous work observed smaller SEP amplitudes in people with stroke with upper limb sensorimotor impairments, compared to age-matched healthy adults (17-19). However, in these studies, a robust clinical assessment of somatosensory function was absent and a different somatosensory stimulus –median nerve stimulation– was applied. We also observed smaller amplitudes of SEP that were only present in persons with stroke and sensorimotor impairments, and not in persons with solely motor impairments. Given the sensory nature of pinprick, this was to be expected (2). However, not in agreement with previous research, the reduced amplitudes of somatosensory evoked potentials were not significantly correlated to the motor clinical score (14). This could be due the different sensory stimuli be applied to the upper limb, as previous work used median nerve stimulation, which involves different ascending sensory pathways and corticospinal projections compared to pinprick, leading to a decreased contribution to the motor impairment (44). However, our results showed the potential of examining peak-to-peak SEP amplitude and beta band resynchronization at baseline for providing information on the magnitude of sensorimotor recovery in the first month poststroke.

Both the beta and alpha frequency bands are involved in sensorimotor processing (45-48). Here, neither type of stroke yielded an impact on alpha or beta band activity following pinpricks, with alpha band activity differences not surviving multiple comparison corrections (p = 0.051). Alpha activity has been linked to cortical inhibitory mechanisms, playing a role in reducing firing in the excitatory pyramidal cells through GABAergic synaptic connections. Attenuated alpha band dynamics in sensorimotor stroke compared to healthy controls may reflect imbalances in recovery mechanisms (49-51). Previous research involving passive movement, revealed that strength of the beta rhythm correlated with hand motor clinical scores post-stroke (23, 52). This partially concurs with our observations, as beta desynchronization was related to motor scores, and beta resynchronization to initial sensorimotor function and change in motor function from T1 to T2. Notably, albeit in the context of aging, previous work from our group also found that beta desynchronization is related to motor performance, with increased desynchronization in healthy older adults being associated to better (inter-limb) coordination (37).

In agreement with our finding, that abnormal delta frequency band power was related to somatosensory impairments, there is accumulating evidence suggesting that low frequency oscillations (LFOs) may be a potential biomarker for upper limb stroke recovery. Activity in the delta frequency band has been identified as a marker in skilled motor control in humans and also in animals (53). People with subacute and chronic stroke with less motor impairment showed a higher delta power in the ipsilesional area (54). Also, longitudinal resting-state EEG shows a decrease in the delta-to-alpha ratio from the subacute to the chronic stage of stroke, and this decrease is associated with a lower National Institute of Health Stroke scale score, i.e., less severity, but not with FMA-UE scores (55).

To our knowledge, our study is the first to examine the neurophysiological mechanisms related to pinprick stimulation in the acute stage post-stroke, both in people with motor- and sensorimotor-impairments. Nevertheless, it is essential to acknowledge the limitations of our exploratory research. While the total sample size of this study, n = 26, was similar to other work in the field, our results focusing on stroke only include a sample of size n = 16 and do not contain longitudinal data, beyond the second visit 7 to 14 days after T1. Therefore, the insights gained from this work in terms of the potential value of pinprick SEPs as a predictor of upper limb sensorimotor recovery in stroke are limited. Furthermore, the pin-prick evoked potential is a mixed potential, implying that both endogenous and exogenous, and therefore also unspecific factors (e.g. attention) may influence it (56). Finally, the varied locations of stroke lesions among our participants hinder the generalizability of our findings.

## Conclusion

Understanding sensorimotor impairment is key for stroke rehabilitation and recovery. Here, the use of several EEG measures proved valuable to identify biomarkers for sensorimotor post-stroke. We observed changes in negative-positive peak-to-peak SEP amplitudes in people with sensorimotor impairments compared to healthy controls. We demonstrated a relationship between beta desynchronization and resynchronization magnitude, and motor and sensorimotor impairment post-stroke. Finally, a larger peak-to-peak SEP amplitude and beta band resynchronization at baseline were related to greater improvements in sensorimotor impairment. Given the exploratory nature of our work, future work should validate these findings in larger sample sizes.

## Acknowledgments

We would like to thank all the participants and their primary caregivers that took part in this research. Additionally, all the clinicians that helped with the recruitment of participants including Ms Adriana Pace and Mrs Noella Sant Cassia from the Physiotherapy Department, Mater Dei Hospital, Malta and Dr André Peeters from the Stroke Unit, Cliniques universitaires Saint-Luc, Brussels, Belgium. Also, Professor Vincent Van Pesch who allowed us to use the EEG equipment in his department at Cliniques universitaires Saint-Luc, Brussels, Belgium. Finally, we would like to thank Professor Stephan Swinnen (KU Leuven) and his team for their EEG expertise and use of their EEG caps.

This work was supported by Promobilia foundation [grant number:17064]. The research work disclosed in this publication is partially funded by the REACH HIGH Scholars Programme – Post-Doctoral Grants awarded to Lisa Tedesco Triccas and an FWO grant (G1129923N) granted to Sybren Van Hoornweder. The project is part-financed by the European Union, Operational Programme II — Cohesion Policy 2014-2020 Investing in human capital to create more opportunities and promote the wellbeing of society -European Social Fund.

The datasets generated and/or analysed during the current study are not publicly available due data protection criteria but are available from the corresponding author on reasonable request.

## Notes

### Competing Interest Statement

The authors have declared no competing interest.

